# Determining organ weight toxicity with Bayesian causal models: Improving on the analysis of relative organ weights

**DOI:** 10.1101/754853

**Authors:** Stanley E. Lazic, Elizaveta Semenova, Dominic P. Williams

**Affiliations:** Data Sciences and Quantitative Biology, AstraZeneca, R&D, Cambridge, CB4 0WG, UK; Functional and Mechanistic Safety, Clinical Pharmacology and Safety Sciences, AstraZeneca, R&D, Cambridge, CB4 0WG, UK

## Abstract

Regulatory authorities require animal toxicity tests for new chemical entities. Organ weight changes are accepted as a sensitive indicator of chemically induced organ damage, but can be difficult to interpret because changes in organ weight might reflect chemically-induced changes in overall body weight. A common solution is to calculate the relative organ weight (organ to body weight ratio), but this inadequately controls for the dependence on body weight – a point made by statisticians for decades, but which has not been widely adopted. The recommended solution is an analysis of covariance (ANCOVA), but it is rarely used, possibly because both the method of statistical correction and the interpretation of the output may be unclear to those with minimal statistical training. Using relative organ weights can easily lead to incorrect conclusions, resulting in poor decisions, wasted resources, and an ethically questionable use of animals. We propose to cast the problem into a causal modelling framework as it directly assesses questions of scientific interest, the results are easy to interpret, and the analysis is simple to perform with freely available software. Furthermore, by taking a Bayesian approach we can model unequal variances, control for multiple testing, and directly provide evidence of safety.

## Introduction

Regulatory authorities require toxicity testing on animals as part of the safety assessment for new drugs, chemicals, biologics, food additives, and medical devices. For new compounds or other chemical entities, organ weight changes are accepted as a sensitive indicator of chemically-induced effects on organs, and are commonly assessed. A problem with interpreting organ weight data is that a compound might affect the overall body weight of animals, and consequently, the size and weight of organs. However, the primary scientific question is whether a compound directly effects an organ, not indirectly through changes in body weight. To overcome this problem, researchers frequently calculate a ratio – called the “relative organ weight” – by dividing each animal’s organ weight by their body weight. The relative organ weights are then plotted, analysed, and interpreted, and researchers assume that this approach will provide correct conclusions. Unfortunately, the ratio fails to properly adjust for differences in body weight between groups – a point made repeatedly but mostly ignored,^1–9^ and which we demonstrate again below.

Michael et al. surveyed organ weight toxicity practices from pharmaceutical, veterinary, chemical, food/nutritional, and consumer product companies in Europe, North America, and Japan^10^. The only response that was consistent across all industries and locations was the use of relative organ weights to adjust for changes in body weight. Michael et al. also reported that respondents “believed that statistical analyses did not always enhance the understanding of these effects and could be misleading.” A remarkable statement, considering that the alternative is an ineffective *ad hoc* adjustment and eye-balling the data, but it nevertheless suggests a dissatisfaction with the recommended statistical procedures. The recommended approach since the 1940s has been the analysis of covariance (ANCOVA)^11^, but it has rarely been used in the past 70 years. Multivariate methods have also been proposed, but have not been adopted either.^12^

In summary, we have a method in widespread use that is known to be ineffective, and an unused but effective method. Publishing yet another paper on the deficiencies of relative organ weights and the merits of ANCOVA is unlikely to change practices. We suspect that many researchers avoid using ANCOVA because they do not understand how it works and how to interpret the output. We therefore propose an alternative approach that is conceptually simple, directly tests the hypothesised causal relationships of interest, and returns results that are intuitive and easy to interpret. We have already used these simple models to show that new neurons in the adult mammalian hippocampus play little or no role in behavioural outcomes, contrary to the accepted view.^13,14^

First, we use simulated data to show how relative organ weights can mislead. Next, we show that an ANCOVA provides the correct conclusion, but is difficult to interpret. Then, we analyse the data with a simple causal model and illustrate how the results are more informative. Finally, we analyse a more complex real data set that has multiple doses and unequal variances across experimental groups for liver weights. We show how the simple causal model can be extended to account for the multiple groups and unequal variances. By taking a Bayesian approach we can make direct probabilistic statements of scientific interest, including the probability of safety (classical frequentist methods can only fail to reject the null hypothesis of “no toxicity”, giving an ambiguous conclusion: is the compound safe, or is power too low to detect an effect?). The Bayesian approach can distinguish between evidence for safety versus “insufficient evidence to make a conclusion”.

## Methods

### Data

The data are from a National Toxicology Program report^15^ and were downloaded from (https://github.com/lahothorn/SiTuR)^16^. Sixty female F344 rats were randomly assigned to the following six groups: 0, 62.5, 125, 250, 500, and 1000 mg of sodium dichromate dihydrate (SDD) per litre of drinking water (10 rats/group). After thirteen weeks, the body and liver weights of the sixty rats were measured, and the question is whether SDD directly affects liver weight. However, since liver weight is related to the overall size of an animal, and SDD affected the body weight of the rats, it is unclear if changes in liver weight merely reflect changes in body weight.

Since the body weight measurements include the weight of the liver, we subtract liver weight from body weight to obtain a “liver-free” estimate of the body weight, which we use in all subsequent analyses. This preprocessing step is rarely mentioned in published studies and therefore presumably not done (or it is unclear if body weights are measured after the relevant organs are removed). Subtracting the liver weight is necessary because otherwise the measured body weight will always reflect changes in the liver weight, although it may make little difference in practice because the liver weight is small compared with the overall body weight. The body and liver weight values were centred by subtracting the mean values (across all dose groups) from each animal’s value. Body and liver weights therefore have a mean of zero, making it easier to define the prior distribution for the group means, but it does not affect the results.

### Simulated data

To illustrate the inadequacy of using relative organ weights, we generated data that has similar means, standard deviations, and correlations as the real data set from the National Toxicology Program report^15^. Because we are generating the data, we know the true causal relationships, and can therefore compare analytical methods to see which provide the correct causal conclusion. Twenty animals per group were simulated with the following equations:

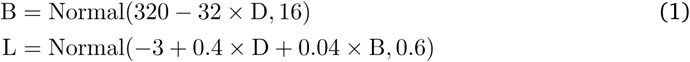

where B is the body weight, L is the liver weight, and D is the dose, which is an indicator variable that equals 0 for control animals and 1 for animals in the drug group.

### Models and analyses

The basic model has three variables: (1) an experimental intervention such as a drug and control group, (2) the weight of a target organ, and (3) body weight (again, this is the remaining body weight after subtracting the liver weight). The scientific hypothesis is whether a difference in organ weight between treatment groups can be attributed to a direct effect of the treatment on the organ, or if the effect is mediated by changes in overall body weight, or both. This basic model is often called a mediation model and it was popularised in the social sciences by Baron and Kenny^17^, and considerable developments have been made since^18–23^. A nice feature of these models is that the hypothesised causal relationships can be expressed as a diagram, where the variables are shown as nodes and the hypothesised causal relationships are indicated with arrows (Fig. 1A).

**Figure 1.**
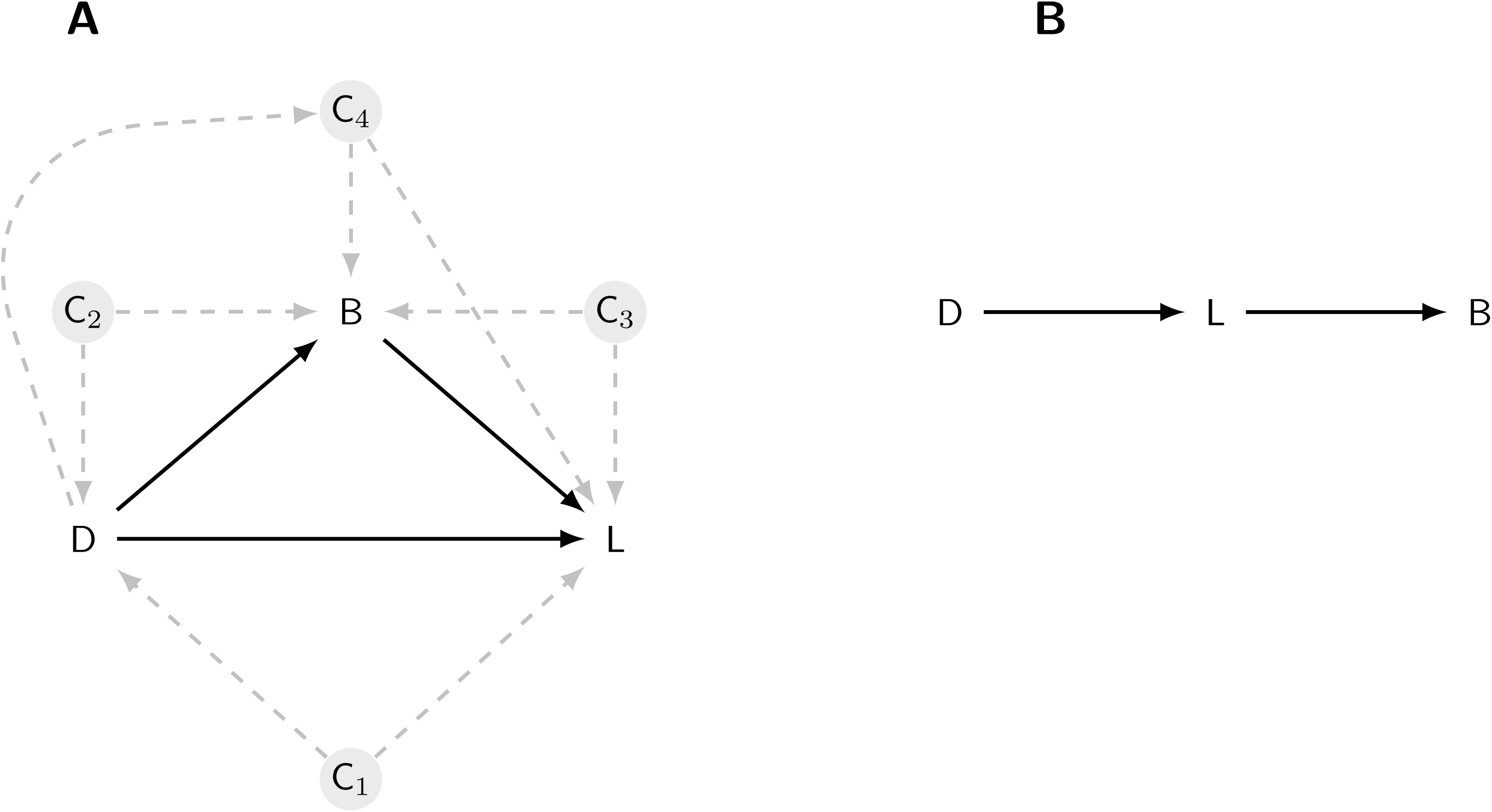
The causal model. A Drug (D) can affect liver weight (L) directly; by altering body weight (B), which in turn affects the liver weight; or through both mechanisms. Four potential confounding effects (C_1_–C_4_) must be ruled out before making causal mechanistic claims about how the drug affects liver weight (A). An alternative causal model is that the drug affects liver weight, making the animal ill, which then leads to a reduction in body weight.

The diagram can then be expressed as a set of equations that are fit to the data. In the equations below we abbreviate body weight to B, liver weight to L, and Dose to D, which can take two levels: D=0 is the control group and D=1 is the treated group. The first equation below describes the effect of the drug on body weight (Drug → Body weight path in Fig. 1A), and the second equation models both the effect of the drug and the effect of body weight on liver weight (Drug → Liver weight and Body weight → Liver weight paths).

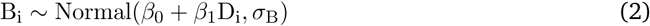

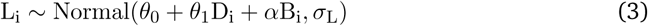

*i* indexes the animal and runs from 1 to the maximum number of animals (*N*). There are seven unknown parameters in these equations (*β*s, *θ*s, *σ*s, and *α*) that are estimated from the data. The three key parameters are: *β*_1_, which quantifies the direction and strength of the drug’s effect on body weight; *α*, which describes how body weight affects liver weight; and *θ*_1_, which quantifies the direction and strength of the drug’s effect on liver weight, after adjusting for the drug’s effect on body weight. Once these parameters are estimated from the data, the following three effects can be calculated. The *direct effect (DE)* of the drug on liver weight (Drug → Liver weight path) is given by *θ*_1_ and is the main hypothesis for this experiment (it corresponds to a standard ANCOVA analysis). The direct effect has an intuitive interpretation as the effect of the drug on liver weight when the body weight is held fixed to what it would have been in the 0 dose group – one can think of it as blocking the Drug → Body weight path in Figure 1A so that the drug can only affect the liver through the direct path. Second, the *indirect effect (IE)* is the product of *β*_1_ and *α* and represents the effect of the drug on the liver weight that is mediated through changes in body weight. Finally, we can calculate the *total effect (TE)* as the sum of the direct and indirect effects. The TE can also be calculated as the difference in the mean liver weights between the two dose groups (ignoring any effect of body weight).

The model has a key assumption about the causal effects: no other variables exist that influence the three variables in the model, which would affect the interpretation of the results. These variables are called confounders and are the nodes C_1_–C_4_ in Figure 1A. The influence of potential confounding variables can be minimised by the design of the experiment when this is feasible, for example by randomising animals to treatment groups. Alternatively, they can be accounted for in the causal model and details of how to incorporate such variables are described in most standard sources; for example, see VanderWeele^23^ for a mathematical description and Tingley et al. for a code-based specification^24^. The first confounder is when a variable affects both organ weight and whether an animal received the control or drug (C_1_ in Fig. 1A). Since this is a randomised experiment, we know how animals were assigned to treatment groups and hence there is no arrow pointing from an unmeasured variable (C_1_) to the Drug node. Contrast this with an observational study comparing people that take aspirin and those that do not. Since people choose to take aspirin (or not) for many reasons (e.g. have headache), there are multiple arrows pointing into the Drug node, and these reasons will also affect the downstream variables of interest. A second confounder is when a variable affects both body weight and whether an animal received the control or drug (C_2_). Again, because this is a randomised experiment, such a variable is unlikely to exist. A third confounder is when a variable affects both liver and body weight (C_3_), and is harder to rule out because we expect that factors such as sex, litter, and age might be relevant. Potential confounders like C_3_ can be dealt with by either holding these values constant experimentally (e.g. using only animals of one sex and all of the same age), or by including these variables as covariates in the mediation model. As we have no further information on the animals in this dataset, we will assume that no such variables exist. A final confounder is when the drug has an effect on a variable (C_4_) that subsequently affects both body weight and organ weight. If such a variable exists, we can still obtain an unbiased estimate of the direct effect, but not the indirect effect. In summary, we can be reasonably certain that the above assumptions about confounding hold in this experiment. The causal modelling approach also requires that hypothesised causal relationships have been properly specified, meaning that the arrows connecting the three variables are pointing in the correct direction. Another potential causal structure is Drug Liver weight Body weight, where the drug has no direct effect on body weight, but has a direct effect on the liver, which makes animals ill, and which subsequently causes them to lose body weight (Fig. 1B). If this is the true relationship, then the analysis is straightforward because we can ignore body weight entirely and examine the liver weights directly (called the Total Effect below). Using relative organ weights in this situation will also bias the results. Fortunately, given that assumptions C_1_–C_4_ hold, the approach described below is valid for both models. If the true relationships are the ones shown in Figure 1A, then the Direct Effect provides the relevant result, but if the true relationships are those shown in Figure 1B, then the Total Effect should be used for inference. We will assume that Figure 1A is the true model of the causal relationships.

The code below shows how to implement the analysis using the `mediation` package in R^24^. The first two lines reflect the two equations above, and the third line takes the output from the first two lines, specifies the treatment variable (`treat = “Dose”`), the mediating variable (`mediator = “B”`), and finally defines the factor levels for the control and treated animals (`control.value = “0”` and `treat.value = “1”`).

~~~
m.mod <- lm(B ∼ Dose)
y.mod <- lm(L ∼ Dose + B)
med.mod <- mediate(m.mod, y.mod, treat = “Dose”, mediator = “B”, control.value
= “0”, treat.value = “1”)
~~~

The above causal model can be extended in two important ways. First, if a confounder like C_3_ in Figure 1B exists, the results could be biased if there is relationship between C_3_ and the treatment conditions; for example, if the control condition has more females and the treated condition has more males^25^. This relationship can result from an imbalance in the design of the experiment and is therefore usually avoidable, but when unavoidable, it can be accounted for by modifying the model definition above (see Tingley et al. for details^24^). We can also test the sensitivity of the conclusions to the assumption that no variable like C_3_ is affecting the results by performing a “what-if” experiment, whereby we induce a positive or negative correlation between body weight and liver weight and recalculate the effects. The `medsens` function in the `mediation` package performs such a sensitivity analysis. Second, the above model assumes that the relationship between liver weight and body weight is constant across treatment groups. Formally, the assumption is that there is no treatment-by-body weight interaction, which is visualised as parallel lines as in Figure 2D. Such an assumption may not hold for a given dataset, and this makes the recommended ANCOVA analysis even more difficult to interpret. With a causal modelling approach such interactions suggest that body weight might *moderate* the effect of the treatment (in addition to mediate); for example, if the drug only affects livers of lighter animals. We do not discuss these extensions here, but see references^17,18,23,26^ for details, and the implementations are described in the documentation for the `mediation` package^24^.

**Figure 2.**
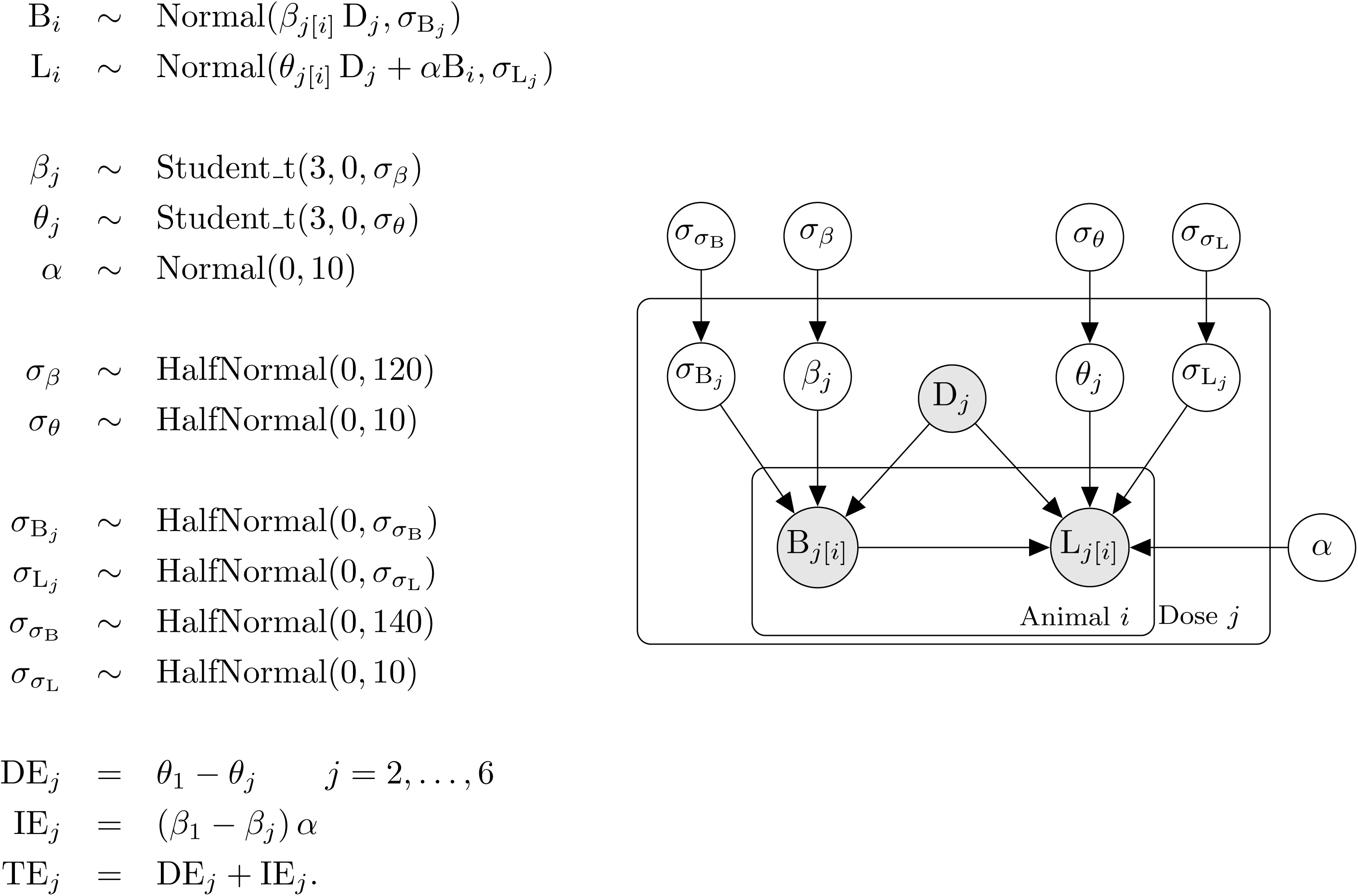
Model definition and representation as a directed acyclic graph (DAG). Symbols are defined in the text.

Although straightforward to implement, the above analysis has several limitations, which we can overcome by switching to a Bayesian analysis that we implement in Stan using the `rstan` package^27,28^. A key benefit of a Bayesian approach is that we obtain straightforward probabilistic statements about quantities of scientific interest, instead of commonly misinterpreted p-values and confidence intervals. Defining the model in Stan enables us to make three extensions. First, we allow for multiple doses or treatment groups and calculate the causal effect at each dose. Second, to control false positives, we shrink the estimates of each group mean towards the overall mean (the mean of all of the data) for both liver weight and body weight. False positives occur when the means of two groups are far apart in a sample of data (relative to the variability of the values within each group), even though there is no true difference between the groups. The traditional way to control false positives is to adjust the p-value upwards (or reduce the threshold for significance) using what has now become a bewildering set of multiple testing corrections. An alternative suggested by Gelman and colleagues is to obtain better estimates of the group means, by using a hierarchical model to shrink the group means towards the global average^29^. In other words, adjust the estimates of the group means instead of adjusting the p-values. Third, we allow the liver weights and body weights to have unequal variances across the dose groups. The assumption of homogeneity of variance is not reasonable for the liver weight values (Fig. 4A). Classical methods either assume a constant variance for each group, which may be inappropriate, or estimate a separate variance for each group using only the data from that group, which can provide poor variance estimates with small sample sizes. The group variances can also be shrunk towards a global average variance, just like the group means, whereby information from all groups is used to estimate the variance for a particular group. This provides more stable estimates while still allowing the variances to differ between groups.

**Figure 4.**
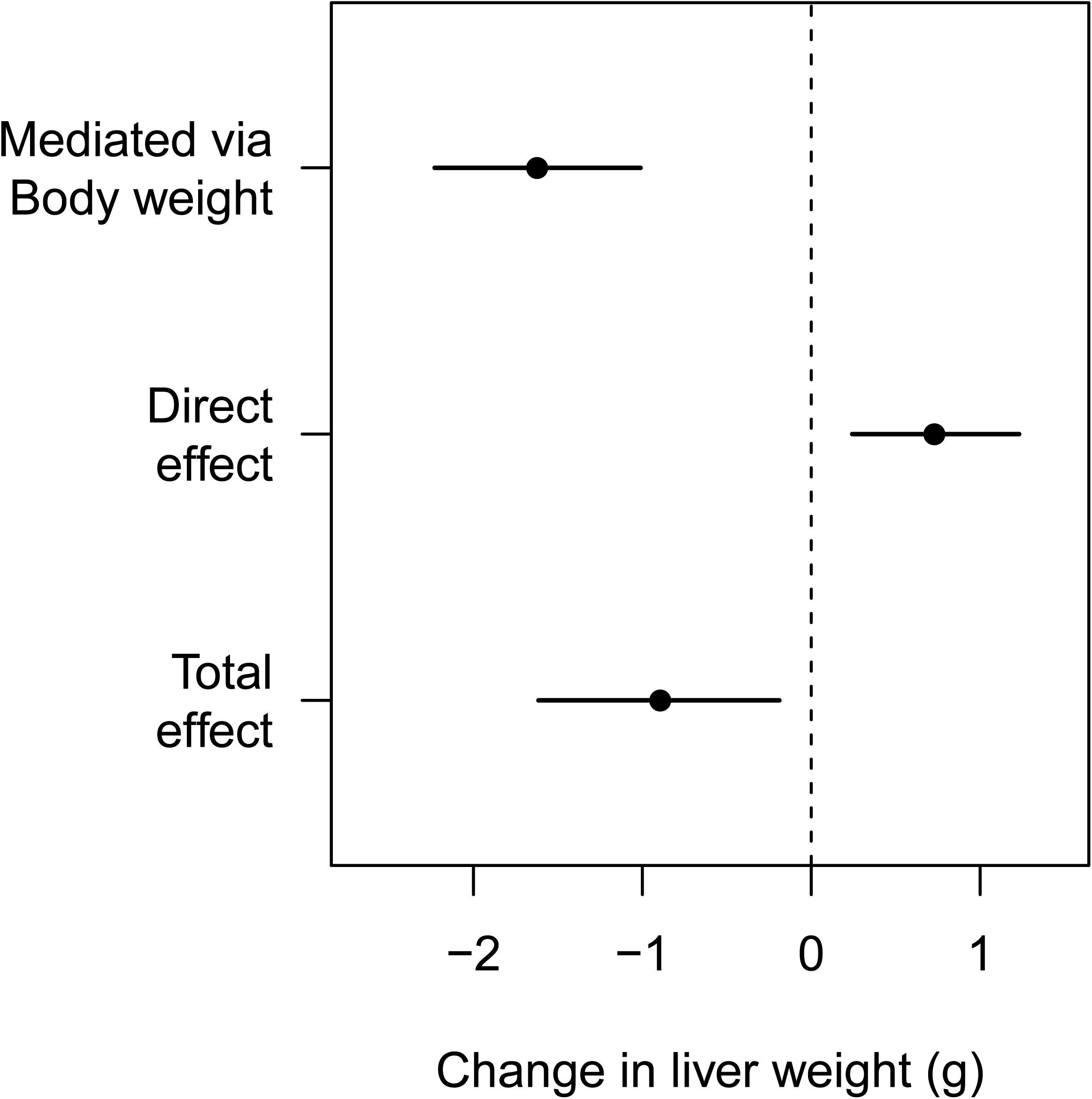
Effect decomposition for the drug’s influence on liver weight. The overall decrease in liver weight is driven by a large decrease in body weight, but the specific and direct effect of the drug is to increase liver weight.

The Bayesian model is defined below and represents our assumptions about the generative process that created the data. It encodes the causal structure from Figure 1A as well as the distributions that describe the liver and body weight values. We do not discuss Bayesian inference in any detail here, but Kruschke and McElreath provide excellent introductions^30,31^. The key idea is that we use distributions (called prior distributions) to represent our knowledge about parameters in the model before observing the current data. The prior distributions are then updated with the data to form posterior distributions, which reflect both the evidence provided by the data and our prior knowledge. All inferences and conclusions are then based on the posterior distributions. In the equations in Figure 2, liver weight, body weight, and the drug variables are once again abbreviated to L, B, and Dose (D), respectively. The model definition is on the left, and graphical notation on the right helps visualise the dependencies.

In the first line of the model (Fig. 2), B is subscripted with the letter *i*, which indexes the animal (*i* runs from 1 to 60). Body weight is modelled as being generated from a normal distribution, and the tilde (*∼*) is read as “is distributed as” or “is generated from” and indicates a stochastic relationship. The distribution of body weights is characterised by a mean (*β*) and standard deviation (*σ*_B_). *j* indexes the dose groups and runs from 1 to 6. Since both *β* and *σ*_B_ are subscripted with a *j*, this indicates that each dose group has its own mean and variance. There are thus six *β* and *σ*_B_ values – one for each group. The *j*[*i*] notation is taken from Gelman and Hill and indicates that animals (*i*) are nested in groups (*j*)^32^. This way of writing the model, where the mean of each group is estimated, is called the “cell means” form (it is more common to include an intercept that represents the mean of the control group, and the other parameters represent the change from the control. The cell means parameterisation is used so that we can more easily define a hierarchical structure to shrink the *β* values towards the global average, and because it makes subsequent calculations easier.) We can examine the *β* values – or differences between *β* values – to determine if differences between groups exist, and hence this first line enables us to test for the effect of the drug on body weight.

In the second line of the model definition, L is liver weight and the *θ* and *σ*_L_ parameters and subscripts have the same interpretation as the *β* and *σ*_B_ parameters in the previous line. *B* is once again the body wight and *α* is the parameter that describes the direction and strength of the relationship between body and liver weight, which is assumed constant across all dose groups, that is, there is no body weight-by-dose interaction.

*β, θ* and *α* are unknown parameters that we estimate from the data. In the Bayesian framework, all unknowns must be represented with a prior distribution, which represents our uncertainty about the unknowns before seeing the data. The next three lines of the model definition specify prior distributions for these unknowns.

Our uncertainty in the six *β* parameters (body weight group means) is represented as a Student-t distribution with three degrees of freedom (df), a mean of zero and a standard deviation of *σ*_*β*_. A t-distribution resembles a normal distribution but has heavier tails, which is controlled by the df (as the df increases a t-distribution becomes a normal distribution). We use a t-distribution because it allows the mean of some dose groups to be far away from others and incorporates our knowledge that large effects may exist at some doses, and because we centred the body weight values, we know that they have a mean of zero. *σ*_*β*_ is the variation between the dose groups – if the drug has no effect then *σ*_*β*_ would be close to zero. The same prior is used for *θ*, which represents the mean liver weight for each group. The prior for *α* is a normal distribution with a mean of zero and a standard deviation of 10.

The next six lines specify priors for the standard deviation parameters in the model. *σ*_*β*_ and *σ*_*θ*_ represent the variation between the group means for body weight and liver weight, respectively. *σ*_B_, and *σ*_L_ represent the variability of the data within each group for body weight and liver weight, and *σ*_*σ*_B and *σ*_*σ*_L represent how much the within group standard deviations (*σ*_B_ and *σ*_L_) vary from group to group for body weight and liver weight. All of the standard deviation parameters in the model are represented with half-normal distributions, which are normal distributions with values less than zero truncated, since standard deviations must be positive. The standard deviations for the halfnormal distributions were set to approximately ten times larger than the empirical values, such that the priors have little influence on the results. The results were similar under reasonable alternative priors (e.g. twice as wide or half as wide).

Finally, given the estimated values for *θ*, the direct effect (DE) of SDD on liver weight can be calculated as the difference of each *θ* from the control group (*θ*_1_). Similarly, the indirect effect (IE) can be calculated as the difference of each *β* from the control group (*β*_1_) times *α*. The total effect (TE) at each dose is just the sum of the direct and indirect effects. The total effect can also be calculated by ignoring body weight and estimating the simple Drug → Liver weight effect. This is not shown in the model specification (Fig. 2) because it is redundant, but it is calculated in the Stan code, represented by the parameter *γ*, as calculating the same quantity in two ways provides a way to check for coding errors.

For comparison, the ANCOVA model is given below, where *θ*_0_ is the intercept and *θ*_1_ – *θ*_4_ are the dose effects. D_1_ – D_4_ are binary dummy variables and the remaining variables and parameters are the same as above. To make this a Bayesian model priors can be placed on the all the parameters.

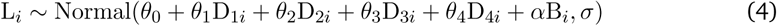

### Controlling false positives

We simulated 1000 datasets with similar properties as the SDD data, including the association between body weight and liver weight, and the within-group variances for both outcomes; but without differences between groups. For each dataset we calculated the direct effect at each dose and deemed the result a false positive if the value of zero was outside of the 95% highest posterior density interval (HPDI). The false positive rate was calculated as the proportion of simulated datasets that had at least one false positive for the direct effect. We provide the data and code to reproduce the analyses in this paper in supplementary file S1.

## Results

### Simulated data

Figure 3A plots the liver weight for twenty control and twenty drug-treated animals, and the drug-treated animals have a reduced liver weight. A classical statistical test for a difference in group means gives a p-value of 0.014. The body weight of the animals in the drug-treated group is also reduced (Fig. 3B; p < 0.001), suggesting that the decreased liver weight might simply reflect changes in body weight. The graph of the relative liver weights (Fig. 3C) and the formal statistical test (p = 0.710) confirms this suspicion, and we might naively conclude that the drug has no direct effect on liver weight. Not only is this conclusion incorrect, but the true effect of the drug is to *increase* liver weight, the opposite of what Figure 3A suggests. Figure 3D shows how this is possible: the drug-treated group (darker points) are shifted to the left of the control group, indicating that their body weight is decreased. The values from the drug-treated group are also shifted above the control group, indicating that the liver weight in the drug-treated group is *increased* for any given value of body weight. Scatterplots and similar graphs can help understand the relationships between liver weight and body weight^33^. Testing for differences in liver weight between groups using the recommended ANCOVA analysis gives a p-value of 0.016, and so we correctly conclude that the drug *increases* liver weight.

**Figure 3.**
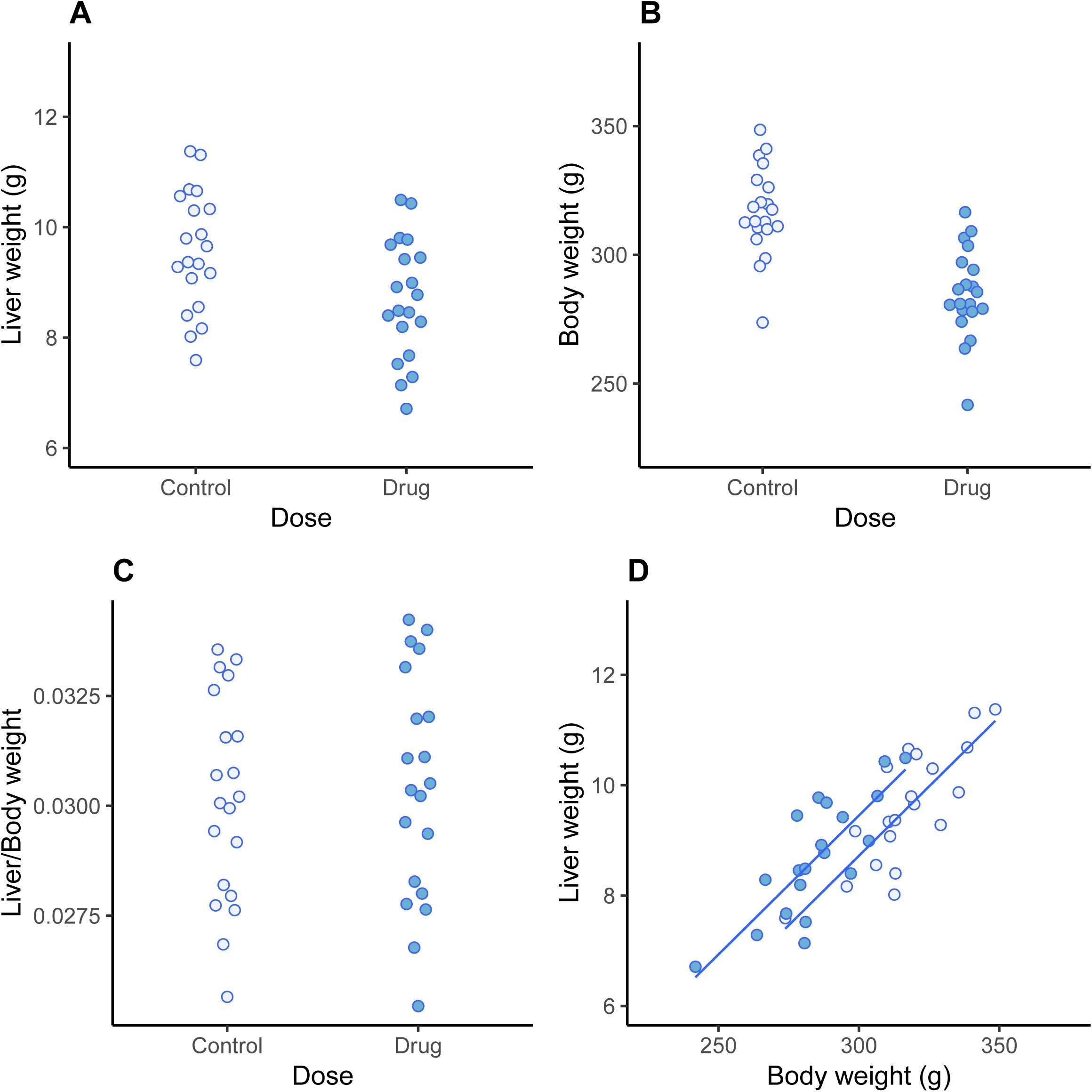
Simulated data. Liver weight is reduced in the drug-treated group (p = 0.014; A), as is body weight (p < 0.001; B). There are no differences between groups for relative organ weights (p = 0.710; C), suggesting (incorrectly) that changes in liver weight simply reflect changes in overall body weight. Relationship between liver and body weight shows that for a given body weight, animals in the drug-treated group have *higher* liver weights than control animals (D).

Figure 4 shows the result of the (non-Bayesian) mediation analysis, which highlights the key effects in the data: (1) the overall liver weight of the animals in the drug-treated group is decreased (total effect), which is driven by (2) a large decrease in body weight, despite (3) the drug directly increasing the liver weight. Two causal effects thus are operating in opposite directions, and which appear to cancel out when looking at the relative liver weight. Liver weight was higher than expected, even after accounting for a decrease in liver weight due to a large decrease in body weight. The sensitivity analysis showed that the conclusion about the indirect effect would not change unless an unknown variable (C_3_) was inducing a correlation of approximately 0.75 between body weight and liver weight. For the direct effect, the conclusion would change if the correlation was 0.25. We can therefore state that only a moderate to large violation of the assumption would change the conclusions. Even though the ANCOVA model came to the correct conclusion, it was not obvious from the model output that two causal effects were working in opposite directions – the raw data needed to be examined along with the model predictions (lines in Fig. 3D).

### Real data

First, we compare group means and standard deviations from a standard non-hierarchical model with those from a hierarchical model. Figure 5 compares the estimated means and 95% Bayesian confidence intervals (highest posterior density intervals) for a standard model that assumes equal variances and a hierarchical model that shrinks the means towards the overall mean (zero in this case because the values were centred) and that allows for unequal variances. Since the group means of the hierarchical model are closer together, the differences between groups is smaller, and hence false positives are less likely.

**Figure 5.**
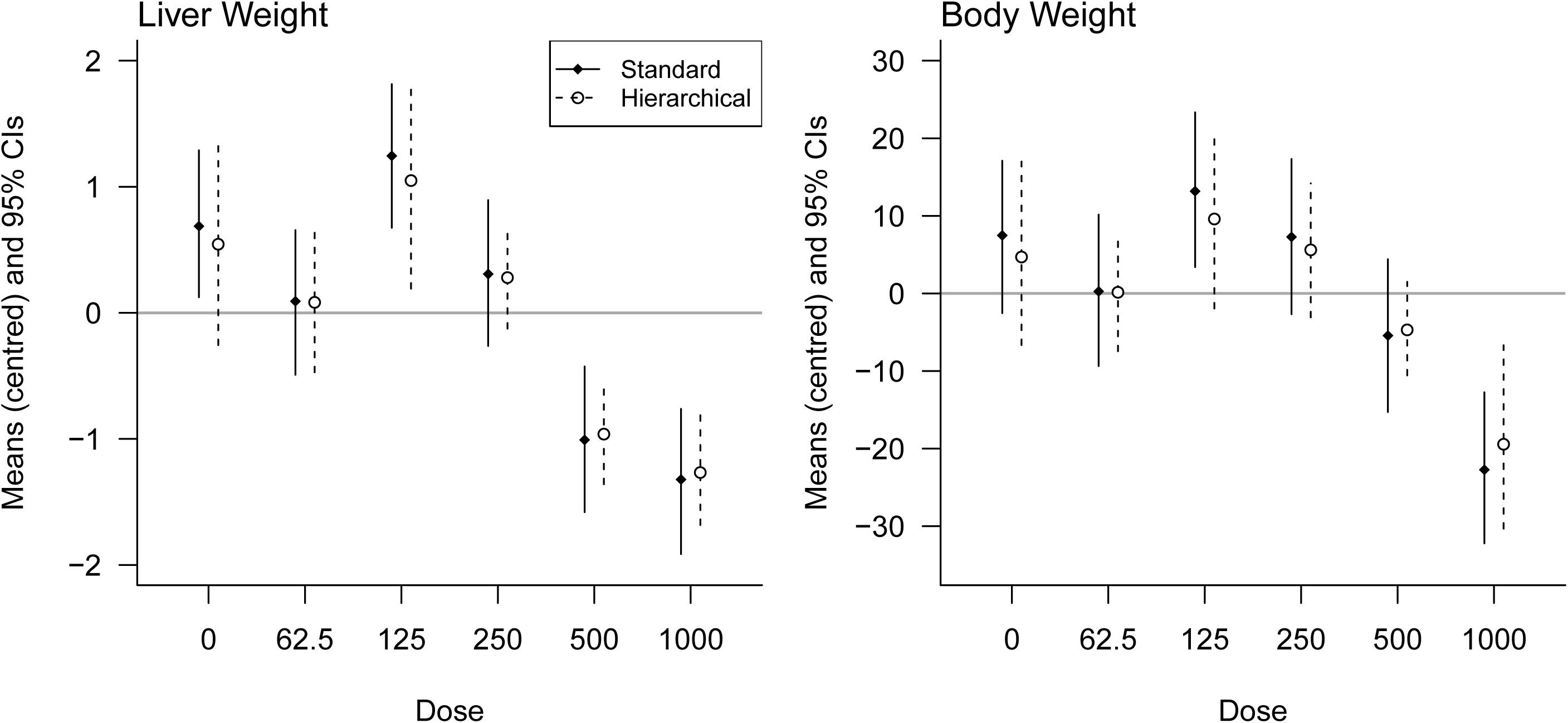
Comparing an equal-variance standard model with a hierarchical model that allows for unequal variances. Note how the hierarchical model shrinks the group means closer to zero (the overall mean), thereby controlling the false positive rate.

For a classical analysis using a 0.05 significance threshold, there is a 23% chance of at least one false positive result when comparing all groups to the control, and a 54% chance when making all possible comparisons. The Bayesian model rephrases hypothesis testing as parameter estimation and “corrections” are not done by changing the p-value or significance threshold but by shrinking the group means towards the overall mean. The simulation experiment showed that when comparing all groups to the control, the false positive rate was only 1% when using the Bayesian model (compared with an expected 23%). Unlike frequentist methods, this approach can control false positives without appreciably increasing false negatives because effects are estimated with greater precision^29^.

Figure 6 shows the raw data from the sodium dichromate dihydrate (SDD) experiment. Qualitatively, both liver weights (A) and body weights (B) decrease at higher doses, as does the liver/body weight ratio (C). As expected, liver and body weights are highly correlated (r = 0.84, p < 0.001; Fig. 6D) but the relative liver weight is still associated with liver weight (r = 0.51, p < 0.001; E), indicating that dividing by body weight provides a poor correction with this data. The recommended ANCOVA method removes the dependence on body weight (r = 0.00, p = 1.00; F). Figure 6A also shows that the liver weight values are less variable at higher doses.

**Figure 6.**
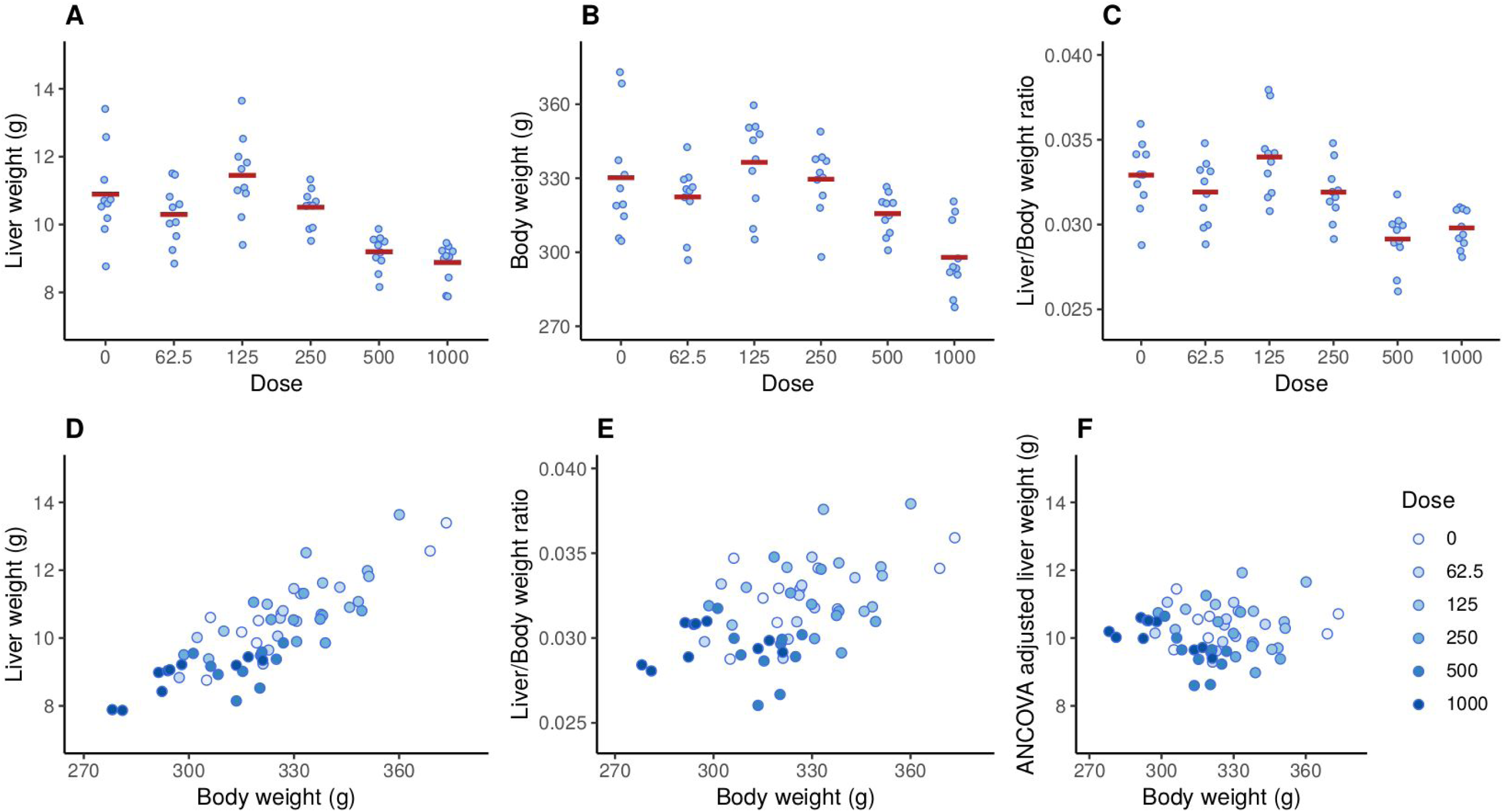
Effect of SDD on liver weight (A), body weight (B), and the liver/body weight ratio (C). Liver and body weight are strongly correlated (r = 0.84, p < 0.001; D), but the relative liver weight is still associated with body weight (r = 0.51, p < 0.001; E), indicating that the dependence on body weight has not been fully removed. The recommend ANCOVA method fully removes the dependence on body weight but can be harder to interpret (F).

Figure 7 shows the results of the Bayesian mediation analysis where each drug-treated group is compared against the control. The left set of graphs are similar to Figure 4, where the total effect of SDD is separated into the direct effect on the liver and the indirect effect that is mediated via a change in body weight. The thick error bars are 50% Bayesian CI and the thin lines are 95% CI. The right set of graphs shows the posterior distribution for the direct effect, which is the main scientific hypothesis. The main summary statistic reported here is the probability that SDD decreases liver weight, indicated as P(Eff < 0). Except for the 125 mg/L group, there is some evidence that SDD decreases liver weight at the lower doses, (P(Eff < 0) = 0.79 and 0.86 at 62.5 and 250 mg/L, respectively), and strong evidence for an effect at higher doses (P(Eff < 0) = 0.99 and 0.97 at 500 and 1000 mg/L). A standard ANCOVA analysis returns the following adjusted p-values (using Holm’s method): at 62.5, 125, and 250 mg/L, p > 0.54; at 500 mg/L p = 0.001; and at 1000 mg/L p = 0.16. The ANCOVA method failed to detect a difference at the highest dose and only the 500 mg/L dose appeared to affect the liver weights. An analysis of the relative liver weights returns the following adjusted p-values (using Holm’s method): at 62.5, 125, and 250 mg/L, p > 0.60; at 500 mg/L p = 0.0002; and at 1000 mg/L p = 0.002. Analysis of the relative liver weights detected a significant difference at both 500 and 1000 doses.

**Figure 7.**
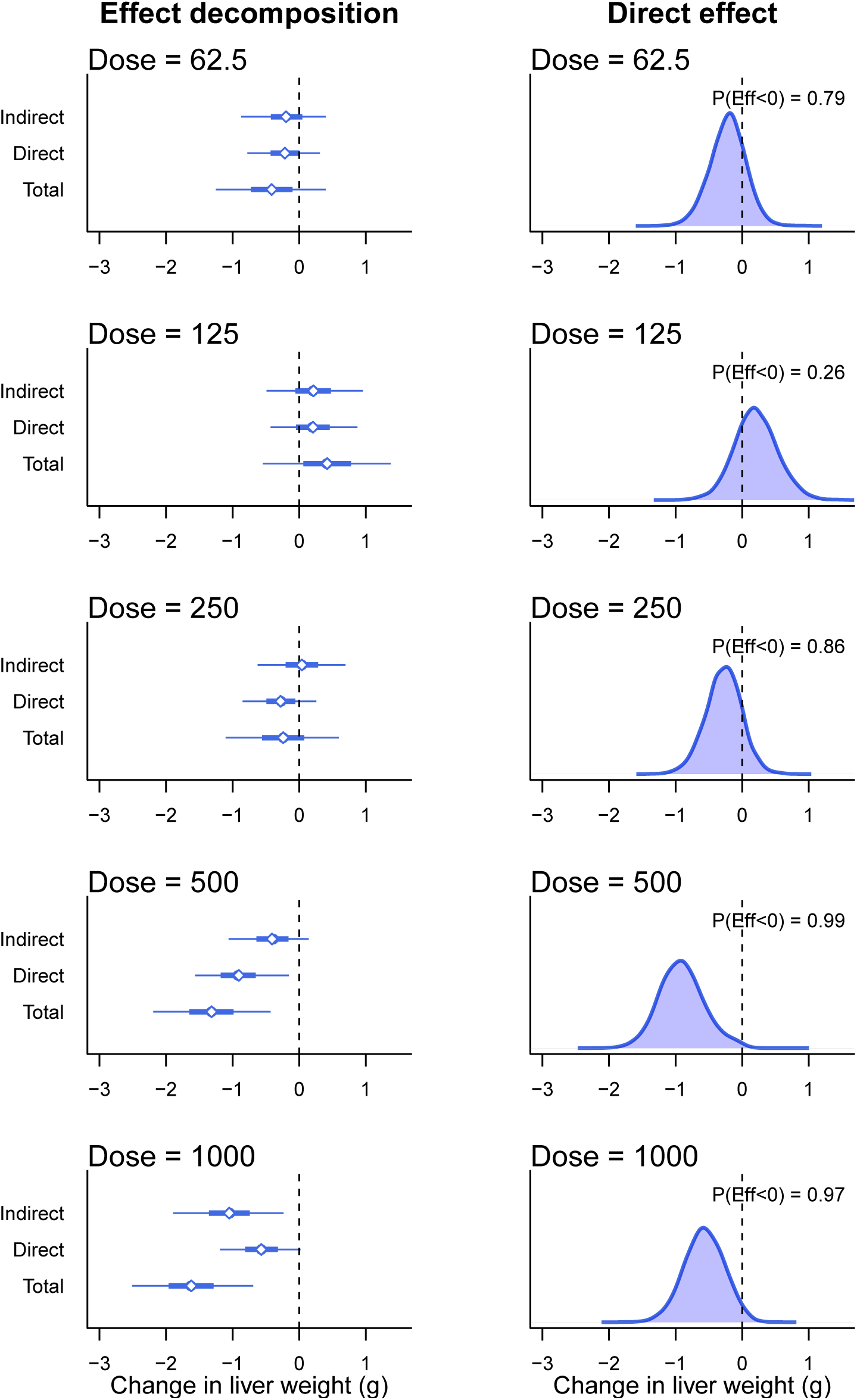
Bayesian mediation analysis. Left graphs show the effect decomposition for the effect of SDD and right graphs show the posterior distribution for the direct effect of SDD on liver weight. P(Eff < 0) is the probability that the effect is less than zero and corresponds to the proportion of the distribution that is below zero.

Instead of detecting the existence of an effect, it is often more important to determine if the change in liver weight is greater than a relevant threshold – usually *±* some value around zero. The threshold can be set such that if the effect exists but is below the threshold, then the effect can be considered negligible. A region of practical equivalence (ROPE) can thus be defined, and if most of the posterior distribution falls within this region, then we can conclude that we have strong evidence of safety^34^. Such a calculation is straightforward and is shown for the 62.5 mg/L dose group in Figure 8. The ROPE is defined as a change of *±* 0.25g for the direct effect (vertical dashed lines) and corresponds to a 20% change in liver weight (a 20% increase in liver weight is the AstraZeneca threshold, whereby bespoke mechanistic investigations into the cause of hepatic hypertrophy or hypercellularity are triggered). Approximately 53% of the posterior distribution for the 62.5 mg/L dose group falls within this region, and we therefore do not have strong evidence of safety at this dose. 44% of the posterior distribution lies below the lower threshold and 4% lies above the upper threshold. A key benefit of this approach is that as the sample size increases, the posterior distribution becomes narrower and will eventually be contained within the ROPE (assuming there is no effect), and we can therefore provide a direct probabilistic statement about safety, which is difficult to do with frequentist methods. For the other doses, the probability that the direct effect is between −0.25 and 0.25 is 0.49, 0.44, 0.04, and 0.15. Thus, at no dose do we have a strong evidence of safety.

**Figure 8.**
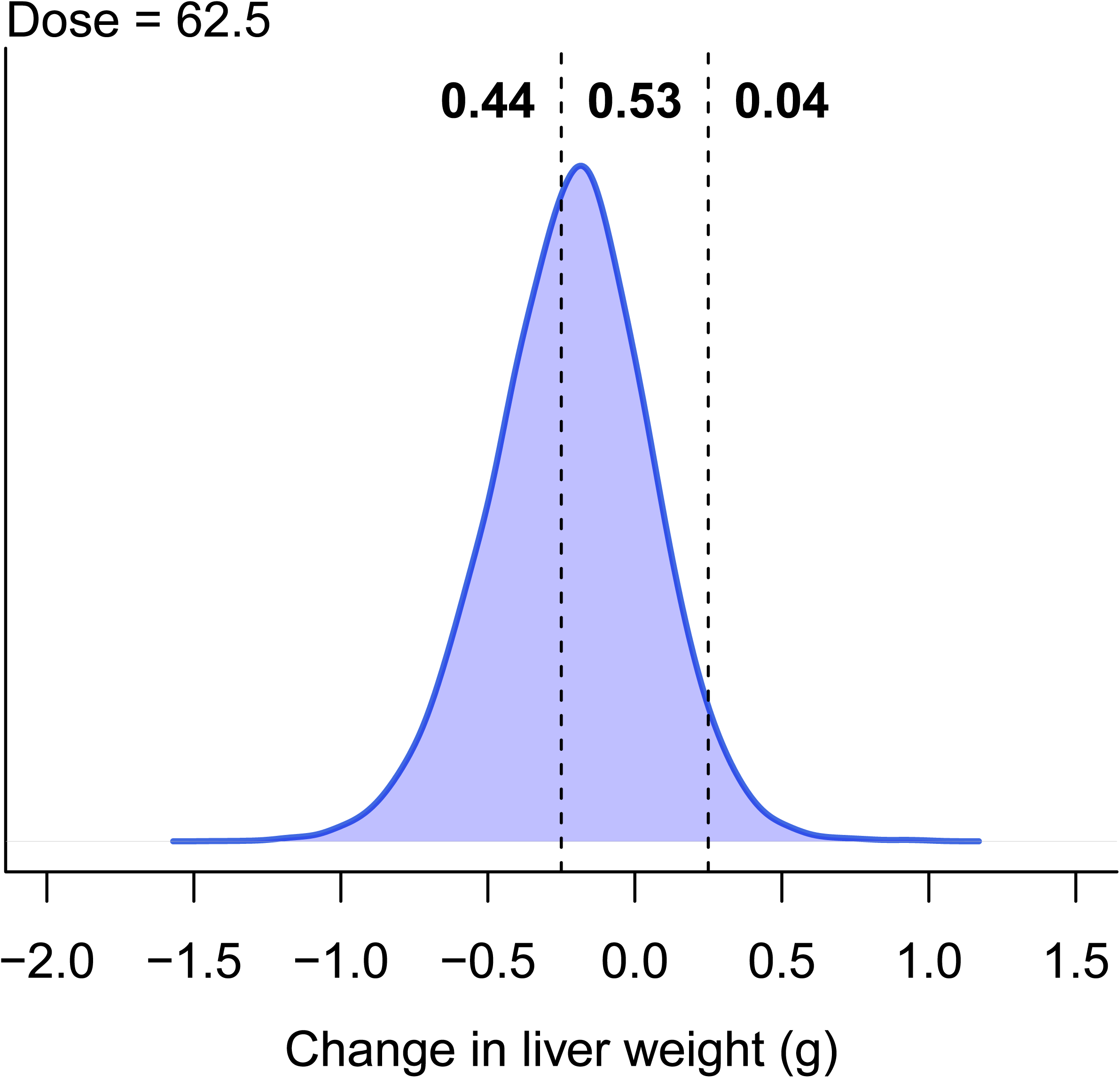
Probability of safety. The vertical lines define the region of practical equivalence, within which the effect is considered negligible. Only 53% of the posterior distribution falls within this region, indicating that there is little evidence of safety.

Another advantage of Bayesian models is the flexibility to easily calculate many quantities of interest. For example, suppose we are interested in the probability that, at any dose, the direct effect is less than a −0.25 g reduction in liver weight. In Figure 8, this is the probability below the left dashed line and equals 0.44. The probability at any dose is the probability at the first dose, or the second dose, or the third dose, and so on, and equals P(DE_62.5_ < −0.25 OR DE_125_ < −0.25 OR DE_250_ < −0.25 OR DE_500_ < −0.25 OR DE_1000_ < −0.25) = 0.98. Hence, even though several doses were tested, we get a single number that enables us to conclude that there is strong evidence of a *biologically relevant* effect of the drug.

## Discussion

The results of the ANCOVA model suggest that SDD does not affect liver weight at the highest dose of 1000 mg/L, based on the adjusted p-value of 0.16 (the unadjusted p-value is 0.04). However, analysing the relative organ weights gives an adjusted p-value of 0.002, suggesting that liver weight *is* affected. Bringing our biological knowledge into play when interpreting the results, we would conclude that if we observe an effect at the second highest dose of 500 mg/L (which all methods found), then it is likely that the effect also exists at a higher dose, especially when the graphs suggest a large effect (Fig. 6A). It appears that the relative organ weight analysis arrived at the correct conclusion, despite its known flaws. Even though the ANCOVA method is preferred over relative organ weights, it may have over-corrected for the changes in body weight in this example. The results of the Bayesian causal model are more consistent with intuition and the relative organ weight results for the highest dose. Figure 7 shows that the *total effect* is larger at 1000 mg/L compared with 500 mg/L, but the indirect effect is a greater proportion of the total effect, thus reducing the magnitude of direct effect. Displaying all of these relationships as in Figure 7 makes the interpretation more straightforward.

In any given experiment, it will be unclear which causal model shown in Figure 1 is correct – assuming these two models exhaust the possibilities. Nevertheless, the Bayesian model described here can estimate unbiased effects under either model. If The Drug → Liver Weight → Body Weight model is correct, then the total effect can be examined. Otherwise the direct effect can be examined. Methods exist to uncover causal structural relationships from data, they are unlikely to be reliable with the small sample sizes used in these toxicology experiments^35^.

A Bayesian model provides direct probabilistic statements about safety by reporting the proportion of the posterior distribution that is within a region of practical equivalence. There was no strong evidence of safety at any dose for the SDD data, but the posterior distributions were all relatively wide, even when the estimated effect was close to zero. This indicates that power was too low to make any strong direct conclusions about safety, although we can still make indirect conclusions for a lack of safety when the posterior is far from zero. This type of analysis opens the possibility of powering experiments by calculating the number of samples needed to achieve confidence intervals of a given width, and not based on the probability of rejecting a null hypothesis as is usually done^36,37^. Powering studies based on the precision of an estimate is also easier than a classic sample size calculation because no minimum effect size needs to be defined.

We showed with simulated data – once again – how relative organ weights can mislead and argue that their use should be discontinued. Graphical displays of the raw values^33^, and mediation models provide a better understanding of the relationships in the data and the effect of chemical substances on organ weights. For simple designs with only two groups the `mediation` R package provides a simple way to fit these models, plot the results, and perform a sensitivity analysis to test the effect of violating assumptions. This analysis can be performed with only a basic knowledge of the free statistical software R, or with JASP’s point-and-click interface (https://jasp-stats.org/), and thus there are no barriers to widespread adoption.

We also showed how the basic model can be extended to account for multiple groups and unequal variances by switching to a Bayesian framework, which also enabled us to control the false positive rate and obtain straightforward probabilistic conclusions about quantities of scientific interest, such as the probability that SDD decreases liver weight or the probability of safety. Although the Bayesian model is more complex and requires knowledge of the Bayesian programming language Stan, it highlights the flexibility of the approach to model the relationships in the data.

We focused on drug-induced organ weight toxicity as a concrete example, but the methods are applicable to many situations where body weight is a confounding factor and where ratio adjustments are currently used. Mediation models can also be used when the mediator, the outcome, or both, are not normally distributed (e.g. binary or count data). These three-variable mediation models are simple examples of larger multi-variables models known as path models or structural equation models, and many areas of experimental biology could benefit from their use^38,39^.

## Acknowledgements

We would like to thank Dr. Felix Kluxen for comments on the manuscript.

## Author contributions

SL: study concept, development statistical model, data analysis, manuscript preparation; ES: data simulation and analysis; DW: study concept, interpretation, manuscript preparation. All authors reviewed the manuscript.

## Data availability

The data used are already publicly available, but have been included in the Supplementary Information for convenience.

