## Supplementary material for "Determining organ weight toxicity with Bayesian causal models: Improving on the analysis of relative organ weights": Data and R code.: 01_standard_mediation_analysis.R

```
library(tidyverse)
library(gridExtra)
library(ggbeeswarm)
library(mediation)

## load data
load("livertox.RData")


## select colours, theme, and panel letter font for plot
cols <- RColorBrewer::brewer.pal(6, "Blues")
theme_set(theme_classic(base_size = 12))
letter <- theme(plot.title = element_text(face="bold"))


## function to simulate organ and body weight data
sim.dat <- function(N, beta0, beta1, theta0, theta1, alpha,
                    sigma.m, sigma.y){

    ## N = Sample size with a group
    ## coefs are defined in the manuscript
    ## sigma.m = SD of mediator (body weight)
    ## sigma.y= SD of outcome (liver weight)
    
    ## groups
    X <- rep(0:1, each=N)

    ## causal model
    M <- beta0 + beta1 * X + rnorm(N, 0, sigma.m)
    Y <- theta0 + theta1 * M + alpha * X + rnorm(N, 0, sigma.y)

    ## calculate relative organ weight
    rel <- Y / M

    return(data.frame(X, M, Y, rel))    
}


## simulate a data set
set.seed(20)
d <- sim.dat(N=20, beta0=320, beta1=-32,
              theta0=-3, theta1=0.04, alpha=0.4, 
              sigma.m=16, sigma.y=0.6)

## results
coef(summary(lm(Y ~ X, data=d)))
coef(summary(lm(Y ~ M + X, data=d)))
coef(summary(lm(M ~ X, data=d)))
coef(summary(lm(rel ~ X, data=d)))


## recode for nicer plots
d <- d %>%
    mutate(cols=recode(X, `0`=cols[1], `1`=cols[4]),
           pch=recode(X, `0`=1, `1`=17),
           X=factor(recode(X, `0`="Control", `1`="Drug")))
    

#pdf("../figs/Sim.pdf", height=7, width=7)
#png("../figs/Sim.png", height=7, width=7, res=300, units="in")
grid.arrange(
    ggplot(d, aes(y=Y, x=X, group=X)) +
    geom_quasirandom(shape=21, color="royalblue", fill=d$cols, cex=1.95, width=0.1) +
    ylab("Liver weight (g)") + xlab("Dose") + ylim(6, 13) + ggtitle("A") +
    letter,

    ggplot(d, aes(y=M, x=X, group=X)) +
    geom_quasirandom(shape=21, color="royalblue", fill=d$cols, cex=1.95, width=0.1) +
    ylab("Body weight (g)") + xlab("Dose") + ylim(225, 375) +
    ggtitle("B") +
    letter,

    ggplot(d, aes(y=rel, x=X, group=X)) +
    geom_quasirandom(shape=21, color="royalblue", fill=d$cols, cex=1.95, width=0.1) +
    ylab("Liver/Body weight") + xlab("Dose") + #ylim(7, 15)
    ggtitle("C") +
    letter,
    
    ggplot(d, aes(y=Y, x=M, group=X)) +
    geom_point(shape=21, color="royalblue", fill=d$cols, cex=1.95) +
    ylab("Liver weight (g)") + xlab("Body weight (g)") + ylim(6, 13) + xlim(225, 375) +
    geom_smooth(method = "lm", se = FALSE, lwd=0.5) + 
    ggtitle("D") + 
    letter,
    
    nrow=2, ncol=2)
#dev.off()


## mediation model
m.mod <- lm(M ~ X, data=d)

## outcome model
y.mod <- lm(Y ~  M + X, data=d)

## fit model
med.mod <- mediate(m.mod, y.mod,
                   treat = "X", mediator = "M",
                   control.value = "Control", treat.value = "Drug",
                   robustSE = TRUE)

summary(med.mod)


##pdf("../figs/mediation_model.pdf", height=4, width=4)
#png("../figs/sim_mediation_model.png", height=4, width=4, res=300, units="in") 
par(las=1,
    mar=c(4.2, 6, 1, 1))
plot(med.mod, xlab="Change in liver weight (g)", yaxt="n")
axis(2, at=1:3, labels = c("Total\neffect", "Direct\neffect", "Mediated via\nBody weight"))
#dev.off()


liv %>%
    mutate(
        ## convert Dose to a factor
        fac.dose=factor(Dose),
        ## ratio adjustment
        ratio=LiverWt/BodyWt,
        ## ANCOVA adjustment
        adj.LiverWt=resid(lm(LiverWt ~ BodyWt)) + mean(LiverWt)) -> 
    liv


## calculate group means for plotting
liv %>%
    group_by(fac.dose) %>% 
    summarise(mean.liver=mean(LiverWt), 
           mean.body=mean(BodyWt),
           mean.ratio=mean(ratio)) ->
    liv.means


#pdf("../figs/Fig1.pdf", height=6, width=11)
#png("../figs/Fig1.png", height=6, width=12, res=300, units="in") 

grid.arrange(
    ggplot(liv, aes(y=LiverWt, x=fac.dose, group=fac.dose)) +
    geom_quasirandom(shape=21, color="royalblue", fill=cols[3], cex=1.2, width=0.1) +
    geom_segment(aes(x = as.numeric(fac.dose)-0.2, y = mean.liver,
                     xend = as.numeric(fac.dose) + 0.2, yend = mean.liver),
                 lwd=1.2, col="firebrick", data=liv.means) + 
    ylab("Liver weight (g)") + xlab("Dose") + ylim(7, 15) + ggtitle("A") +
    letter, 

    ggplot(liv, aes(y=BodyWt, x=fac.dose, group=fac.dose)) +
    geom_quasirandom(shape=21, color="royalblue", fill=cols[3], cex=1.2, width=0.1) +
    geom_segment(aes(x = as.numeric(fac.dose)-0.2, y = mean.body,
                     xend = as.numeric(fac.dose) + 0.2, yend = mean.body),
                 lwd=1.2, col="firebrick", data=liv.means) + 
    ylab("Body weight (g)") + xlab("Dose") + ylim(270,380) + ggtitle("B") +
    letter, 

    ggplot(liv, aes(y=ratio, x=fac.dose, group=fac.dose)) +
    geom_quasirandom(shape=21, color="royalblue", fill=cols[3], cex=1.2, width=0.1) +
    geom_segment(aes(x = as.numeric(fac.dose)-0.2, y = mean.ratio,
                     xend = as.numeric(fac.dose) + 0.2, yend = mean.ratio),
                 lwd=1.2, col="firebrick", data=liv.means) + 
    ylab("Liver/Body weight ratio") + xlab("Dose") + ylim(0.024,0.04) + ggtitle("C") +
    letter, 

    ggplot(liv, aes(y=LiverWt, x=BodyWt, fill=fac.dose)) + 
    geom_point(shape=21, color="royalblue", size=2) +
    scale_fill_manual(name = "Dose", values=cols) +
    ylab("Liver weight (g)") +  xlab("Body weight (g)") +
    xlim(270,380) + ylim(7, 15) + ggtitle("D") +
    letter + theme(legend.position = "none"), 

    ggplot(liv, aes(y=ratio, x=BodyWt, fill=fac.dose)) + 
    geom_point(shape=21, color="royalblue", size=2) +
    scale_fill_manual(name = "Dose", values=cols) +
    ylab("Liver/Body weight ratio") +  xlab("Body weight (g)") + ggtitle("E") +
    ylim(0.024, 0.04) + xlim(270, 380) +
    letter + theme(legend.position = "none"), 
    
    ggplot(liv, aes(y=adj.LiverWt, x=BodyWt, fill=fac.dose)) + 
    geom_point(shape=21, color="royalblue", size=2) +
    scale_fill_manual(name = "Dose", values=cols) +
    ylab("ANCOVA adjusted liver weight (g)") +
    xlab("Body weight (g)") + ylim(7, 15) + xlim(270,380) + ggtitle("F") +
    letter, 

    nrow=2, ncol=3)
#dev.off()


## check how adjustments work
cor.test(~LiverWt + BodyWt, data=liv)
cor.test(~ratio + BodyWt, data=liv)
cor.test(~adj.LiverWt + BodyWt, data=liv)


## analysis of ratios
liv.mod <- lm(LiverWt ~ fac.dose, data=liv)
summary(liv.mod)
par(mfrow=c(2, 2))  
plot(liv.mod)


## analysis of ratios
rel.mod <- lm(ratio ~ fac.dose, data=liv)
summary(rel.mod)
par(mfrow=c(2, 2))
plot(rel.mod)

## calculate 95% CI
cis <- confint(rel.mod)[-1, ]

plot(c(1:5) ~ coef(rel.mod)[-1], xlim=c(-0.006, 0.006))
segments(cis[, 1], 1:5, cis[, 2], 1:5)
abline(v=0, lty=2)
```
