## Supplementary material for "Determining organ weight toxicity with Bayesian causal models: Improving on the analysis of relative organ weights": Data and R code.: 02_Bayesian_models.R

```
library(tidyverse)
library(mcmcplots)
library(rstan)
library(rstanarm)
library(gridExtra)
library(extrafont)

source("functions.R")

options(mc.cores = parallel::detectCores())
rstan_options(auto_write = TRUE)


## load data
load("livertox.RData")


liv %>%
    mutate(
        ## convert Dose to a factor
        fac.dose = factor(Dose),
        ## calc liver-free body weight
        bw = BodyWt - LiverWt, 
        ## center body and liver weight (helps to set the prior)
        c.BodyWt = bw - mean(bw),
        c.LiverWt = LiverWt - mean(LiverWt)) -> 
    liv


## get an idea of values to help set priors
tapply(liv$c.BodyWt, liv$fac.dose, mean)
mean(tapply(liv$c.BodyWt, liv$fac.dose, sd)) # sigma
sd(tapply(liv$c.BodyWt, liv$fac.dose, mean)) # grp_sigma

tapply(liv$c.LiverWt, liv$fac.dose, mean)
mean(tapply(liv$c.LiverWt, liv$fac.dose, sd)) # sigma
sd(tapply(liv$c.LiverWt, liv$fac.dose, mean)) # grp_sigma


## Standard model with equal variances
std.bw <- stan_glm(c.BodyWt ~ 0 + fac.dose, data=liv, iter=10000,
                    prior_intercept = normal(0, 50, autoscale=FALSE),
                    prior = normal(0, 120, autoscale=FALSE),
                    prior_aux = normal(0, 140, autoscale=FALSE))

std.lw <- stan_glm(c.LiverWt ~ 0 + fac.dose, data=liv, iter=10000,
                  prior_intercept = normal(10, 20, autoscale=FALSE),
                  prior = normal(0, 10, autoscale=FALSE),
                  prior_aux = normal(0, 10, autoscale=FALSE))

## extract draws from posterior
post.std.bw <- as.data.frame(std.bw)
post.std.lw <- as.data.frame(std.lw)

## calculate means and 95% HPDI for plotting
post.std.bw %>%
    select_at(vars(fac.dose0:fac.dose1000)) %>%
    calc.stats() ->
    std.bw.result

post.std.lw %>%
    select_at(vars(fac.dose0:fac.dose1000)) %>%
    calc.stats() ->
    std.lw.result


    

## Hierarchical model with unequal variances
compiled.model <- stan_model("multilev_het_var.stan")

## data to feed into model
d.hier <- with(liv,
               list(
                   N = nrow(liv),
                   P = length(unique(fac.dose)),
                   grp = as.integer(as.numeric(fac.dose)),
                   bw = c.BodyWt,
                   lw = c.LiverWt,
                   prior_only = 0)
               )

## draw samples from posterior
fit.hier <- sampling(compiled.model, data=d.hier, iter=10000, chains=3,
                      seed=123)

check_hmc_diagnostics(fit.hier)

## get draws from posterior
post.hier <- rstan::extract(fit.hier)


## calculate means and 95% HPDI for plotting
post.hier$beta %>%
    calc.stats() ->
    hier.bw.result

post.hier$gamma %>%
    calc.stats() ->
    hier.lw.result


#png("../figs/shrinkage.png", height=4.25, width=9, res=300, units="in") 
par(las=1,
    mfrow=c(1, 2),
    mar=c(4, 4, 2, 1))
plot.new()

plot.window(ylim=c(-2, 2), xlim=c(0.5, 6.5))
abline(h=0, col="darkgrey", lwd=1.5)

with(std.lw.result, segments(I(c(1:6) - 0.1), lower, I(c(1:6) - 0.1), upper))
with(hier.lw.result, segments(I(c(1:6) + 0.1), lower, I(c(1:6) + 0.1), upper, lty=2))

points(means ~ I(c(1:6) - 0.1), data=std.lw.result, pch=18, cex=0.8)
points(means ~ I(c(1:6) + 0.1), data=hier.lw.result, pch=21, bg="white", cex=0.8)

axis(1, at=1:6, labels=std.lw.result$dose) ; axis(2) ; box(bty="L")
mtext("Liver Weight", cex=1.2, adj=0, line=0.2)
mtext("Dose", side=1, line=3)
par(las=0)
mtext("Means (centred) and 95% CIs", side=2, line=3)

legend("topright", lty=1:2, pch=c(18, 21),
       legend = c("Standard", "Hierarchical"), cex=0.8)


par(las=1)
plot.new()
plot.window(ylim=c(-35, 30), xlim=c(0.5, 6.5))
abline(h=0, col="darkgrey", lwd=1.5)

with(std.bw.result, segments(I(c(1:6) - 0.1), lower, I(c(1:6) - 0.1), upper))
with(hier.bw.result, segments(I(c(1:6) + 0.1), lower, I(c(1:6) + 0.1), upper, lty=2))

points(means ~ I(c(1:6) - 0.1), data=std.bw.result, pch=18, cex=0.8)
points(means ~ I(c(1:6) + 0.1), data=hier.bw.result, pch=21, bg="white", cex=0.8)

axis(1, at=1:6, labels=std.bw.result$dose) ; axis(2) ; box(bty="L")
mtext("Body Weight", cex=1.2, adj=0, line=0.2)
mtext("Dose", side=1, line=3)
par(las=0)
mtext("Means (centred) and 95% CIs", side=2, line=3)

#dev.off()


## plot coefs
plot(fit.hier, par="beta")


## calculate effects for each dose
D1 <- calc.effects(post.hier$theta[, 2] - post.hier$theta[, 1],
                   post.hier$beta[, 2] - post.hier$beta[, 1],
                   post.hier$alpha)
D2 <- calc.effects(post.hier$theta[, 3] - post.hier$theta[, 1],
                   post.hier$beta[, 3] - post.hier$beta[, 1],
                   post.hier$alpha)
D3 <- calc.effects(post.hier$theta[, 4] - post.hier$theta[, 1],
                   post.hier$beta[, 4] - post.hier$beta[, 1],
                   post.hier$alpha)
D4 <- calc.effects(post.hier$theta[, 5] - post.hier$theta[, 1],
                   post.hier$beta[, 5] - post.hier$beta[, 1],
                   post.hier$alpha)
D5 <- calc.effects(post.hier$theta[, 6] - post.hier$theta[, 1],
                   post.hier$beta[, 6] - post.hier$beta[, 1],
                   post.hier$alpha)
                                      

## density plots for the direct effects
dens.D1 <- density(D1$Direct, adjust=1.5)
dens.D2 <- density(D2$Direct, adjust=1.5)
dens.D3 <- density(D3$Direct, adjust=1.5)
dens.D4 <- density(D4$Direct, adjust=1.5)
dens.D5 <- density(D5$Direct, adjust=1.5)


#png("../figs/posteriors.png", height=8, width=5, res=300, units="in") 
par(mfrow=c(5,2),
    lend="square",
    mar=c(3,5,2,0.1),
    oma=c(0,0,2,0),
    font.axis=1)

caterplot(D1, style="plain", col="royalblue", pch=16, reorder=FALSE, 
          quantiles=list(outer=c(0.025,0.975),inner=c(0.25,0.75)),
          val.lim=c(-3, 1.5), cex.labels=1, lwd=c(1,3), bty="L")
abline(v=0, lty=2)
points(c(3:1) ~ apply(D1, 2, mean), pch=23, bg="white", col="royalblue")
mtext("Dose = 62.5", side=3, line=0, cex=1, adj=0)
mtext("Effect decomposition", side=3, line=2, cex=1, font=2)


plot(dens.D1, lwd=2, col="royalblue", ylab="", bty="L",
     xlab="", yaxt="n", main="", xlim=c(-3, 1.5), ylim=c(0, 2))
polygon(dens.D1$x, dens.D1$y, col=rgb(0, 0, 1, 0.25), border=NA)
abline(v=0, lty=2)
mtext("Dose = 62.5", side=3, line=0, cex=1, adj=0)
legend("topright", bty = "n",
       legend=paste0("P(Eff<0) = ", round(mean(D1$Direct < 0), 2)))
mtext("Direct effect", side=3, line=2, cex=1, font=2)

caterplot(D2, style="plain", col="royalblue", pch=19, reorder=FALSE, 
          quantiles=list(outer=c(0.025,0.975),inner=c(0.25,0.75)),
           val.lim=c(-3, 1.5), cex.labels=1, lwd=c(1,3), bty="L")
abline(v=0, lty=2)
points(c(3:1) ~ apply(D2, 2, mean), pch=23, bg="white", col="royalblue")
mtext("Dose = 125", side=3, line=0, cex=1, adj=0)

plot(dens.D2, lwd=2, col="royalblue", ylab="", bty="L",
     xlab="", yaxt="n", main="", xlim=c(-3, 1.5), ylim=c(0, 2))
polygon(dens.D2$x, dens.D2$y, col=rgb(0, 0, 1, 0.25), border=NA)
abline(v=0, lty=2)
mtext("Dose = 125", side=3, line=0, cex=1, adj=0)
legend("topright", bty = "n",
       legend=paste0("P(Eff<0) = ", round(mean(D2$Direct < 0), 2)))


caterplot(D3, style="plain", col="royalblue", pch=19, reorder=FALSE, 
          quantiles=list(outer=c(0.025,0.975),inner=c(0.25,0.75)),
           val.lim=c(-3, 1.5), cex.labels=1, lwd=c(1,3), bty="L")
abline(v=0, lty=2)
points(c(3:1) ~ apply(D3, 2, mean), pch=23, bg="white", col="royalblue")
mtext("Dose = 250", side=3, line=0, cex=1, adj=0)

plot(dens.D3, lwd=2, col="royalblue", ylab="", bty="L",
     xlab="", yaxt="n", main="", xlim=c(-3, 1.5), ylim=c(0, 2))
polygon(dens.D3$x, dens.D3$y, col=rgb(0, 0, 1, 0.25), border=NA)
abline(v=0, lty=2)
mtext("Dose = 250", side=3, line=0, cex=1, adj=0)
legend("topright", bty = "n",
       legend=paste0("P(Eff<0) = ", round(mean(D3$Direct < 0), 2)))


caterplot(D4, style="plain", col="royalblue", pch=19, reorder=FALSE, 
          quantiles=list(outer=c(0.025,0.975),inner=c(0.25,0.75)),
           val.lim=c(-3, 1.5), cex.labels=1, lwd=c(1,3), bty="L")
abline(v=0, lty=2)
points(c(3:1) ~ apply(D4, 2, mean), pch=23, bg="white", col="royalblue")
mtext("Dose = 500", side=3, line=0, cex=1, adj=0)

plot(dens.D4, lwd=2, col="royalblue", ylab="", bty="L",
     xlab="", yaxt="n", main="", xlim=c(-3, 1.5), ylim=c(0, 2))
polygon(dens.D4$x, dens.D4$y, col=rgb(0, 0, 1, 0.25), border=NA)
abline(v=0, lty=2)
mtext("Dose = 500", side=3, line=0, cex=1, adj=0)
legend("topright", bty = "n",
       legend=paste0("P(Eff<0) = ", round(mean(D4$Direct < 0), 2)))


caterplot(D5, style="plain", col="royalblue", pch=19, reorder=FALSE, 
          quantiles=list(outer=c(0.025,0.975),inner=c(0.25,0.75)),
           val.lim=c(-3, 1.5), cex.labels=1, lwd=c(1,3), bty="L")
abline(v=0, lty=2)
points(c(3:1) ~ apply(D5, 2, mean), pch=23, bg="white", col="royalblue")
mtext("Dose = 1000", side=3, line=0, cex=1, adj=0)
mtext("Change in liver weight (g)", side=1, line=2, cex=0.8, font=1)

plot(dens.D5, lwd=2, col="royalblue", ylab="", bty="L",
     xlab="", yaxt="n", main="", xlim=c(-3, 1.5), ylim=c(0, 2))
polygon(dens.D5$x, dens.D5$y, col=rgb(0, 0, 1, 0.25), border=NA)
mtext("Change in liver weight (g)", side=1, line=2, cex=0.8, font=1)
abline(v=0, lty=2)
mtext("Dose = 1000", side=3, line=0, cex=1, adj=0)
legend("topright", bty = "n",
       legend=paste0("P(Eff<0) = ", round(mean(D5$Direct < 0), 2)))

#dev.off()


## prob of safety
tapply(liv$LiverWt, liv$fac.dose, mean)

## interested in a 20% effect
(10.894 * 0.2) # +/- 2.1788 g

## proportion of posterior between +/- 20%
mean(D1$Direct > -0.25 & D1$Direct < 0.25)
mean(D2$Direct > -0.25 & D2$Direct < 0.25)
mean(D3$Direct > -0.25 & D3$Direct < 0.25)
mean(D4$Direct > -0.25 & D4$Direct < 0.25)
mean(D5$Direct > -0.25 & D5$Direct < 0.25)

## proportion of posterior below and above threshold
mean(D1$Direct < -0.25)
mean(D1$Direct > 0.25)


#png("../figs/ROPE.png", height=4, width=4, res=300, units="in") 
par(mar=c(4, 1, 2, 0.1))

plot(dens.D1, lwd=2, col="royalblue", ylab="", bty="L",
     xlab="", yaxt="n", main="", xlim=c(-2, 1.5), ylim=c(0, 1.75))
polygon(dens.D1$x, dens.D1$y, col=rgb(0, 0, 1, 0.25), border=NA)
abline(v=c(-0.25, 0.25), lty=2)
mtext("Dose = 62.5", side=3, line=0, cex=1, adj=0)
mtext("Change in liver weight (g)", side=1, line=2.5)

text(y=1.7, x=0, label="0.53", font=2)
text(y=1.7, x=0.5, label="0.04", font=2)
text(y=1.7, x= -0.5, label="0.44", font=2)

#dev.off()


## prob that effect is < -0.25 for each dose
mean(D1$Direct < -0.25)
mean(D2$Direct < -0.25)
mean(D3$Direct < -0.25)
mean(D4$Direct < -0.25)
mean(D5$Direct < -0.25)


# prob that at least one dose has an effect < -0.25
mean(D1$Direct < -0.25 |
     D2$Direct < -0.25 |
     D3$Direct < -0.25 |
     D4$Direct < -0.25 |
     D5$Direct < -0.25)
```
