## Supplementary material for "Determining organ weight toxicity with Bayesian causal models: Improving on the analysis of relative organ weights": Data and R code.: 03_simulations_false_positives.R

```
library(rstan)
library(tidyverse)
library(mcmcplots)

options(mc.cores = parallel::detectCores())
rstan_options(auto_write = TRUE)

source("functions.R")

## compile model for simulating data
comp <- stan_model("simulations.stan")

## simulate data
Nsets = 1000 # number of simulated data sets
sim.dat <- sampling(comp, data=list(N=60), iter=Nsets, warmup=0, chains=1,
                      seed=123, algorithm = "Fixed_param")
  
post.sim.dat <- rstan::extract(sim.dat)

## Hierarchical model with unequal variances
compiled.model <- stan_model("multilev_het_var.stan")

## make a directory to store results
system("mkdir fits")


## fit model to each simulated data set
for (i in 1:Nsets){
  
  d.hier <-   list(  N = 60,
                     P = 6,
                     grp = rep(1:6, each=10),
                     bw = post.sim.dat$bw_sim[i, ],
                     lw = post.sim.dat$lw_sim[i, ],
                     prior_only = 0)
  
  ## draw samples from posterior
  fit.hier <- sampling(compiled.model, data=d.hier, iter=8000, chains=3,
                       seed=123, control=list(adapt_delta=0.9))
  
  save(fit.hier, file=paste0("fits/", i, ".Rda"))
}


## vector to store results of direct effect
results = rep(NA, Nsets)

for (i in 1:Nsets){
  load(paste0("fits/",i,".Rda"))
  
  post.hier <- rstan::extract(fit.hier)
  
  ## calculate effects for each dose
  D1 <- calc.effects(post.hier$theta[, 2] - post.hier$theta[, 1],
                     post.hier$beta[, 2] - post.hier$beta[, 1],
                     post.hier$alpha)
  D2 <- calc.effects(post.hier$theta[, 3] - post.hier$theta[, 1],
                     post.hier$beta[, 3] - post.hier$beta[, 1],
                     post.hier$alpha)
  D3 <- calc.effects(post.hier$theta[, 4] - post.hier$theta[, 1],
                     post.hier$beta[, 4] - post.hier$beta[, 1],
                     post.hier$alpha)
  D4 <- calc.effects(post.hier$theta[, 5] - post.hier$theta[, 1],
                     post.hier$beta[, 5] - post.hier$beta[, 1],
                     post.hier$alpha)
  D5 <- calc.effects(post.hier$theta[, 6] - post.hier$theta[, 1],
                     post.hier$beta[, 6] - post.hier$beta[, 1],
                     post.hier$alpha)
  
  # number of estimated direct effects that fall outside of the 95% HPDI
  DE.df <- cbind(D1$Direct, 
                 D2$Direct,
                 D3$Direct,
                 D4$Direct,
                 D5$Direct)
  
  means <- colMeans(DE.df)
  cis <- matrix(as.numeric(HPDinterval(mcmc(DE.df))), ncol = 2)
  mcis <- data.frame(cbind(means, cis))
  
  mcis$outside  <-  rep(NA, nrow(mcis))

  for (r in 1:nrow(mcis)){
      ## lower CI above zero or upper CI below zero
      mcis$outside[r] = mcis[r, 2] >= 0  |  mcis[r, 3] <= 0
  }
  
  results[i]  <-  any(mcis$outside) # at least 1 false positive?
  
  print(results[i])
}

round(mean(results, na.rm=TRUE), 4)
```
