## Supplementary material for "Determining organ weight toxicity with Bayesian causal models: Improving on the analysis of relative organ weights": Data and R code.: functions.R

```
## calculate effect decompositions
calc.effects <- function(theta, beta, alpha){

    ## Direct effect
    DE <- theta

    ## Mediated (indirect) effect
    IE <- beta * alpha
    
    ## total effect
    TE <- IE + DE

    df <- data.frame(Indirect=IE, Direct=DE, Total=TE)
    return(df)
}


##  calculates summary statistics of posterior distribution
calc.stats <- function(x){
    means <- colMeans(x)
    cis <- HPDinterval(mcmc(x))
    return(data.frame(means, cis, dose=factor(c(0, 62.5, 125, 250, 500, 1000))))
}
```
