## Supplementary material for "Determining organ weight toxicity with Bayesian causal models: Improving on the analysis of relative organ weights": Data and R code.: multilev_het_var.stan

```
data{
  int N;                                // Number of animals
  int P;                                // Number of groups
  int grp[N];                           // Group indicator
  vector[N] bw;                         // Body weights
  vector[N] lw;                         // Liver weights
  int<lower = 0, upper = 1> prior_only; // Sample from prior only?
}

parameters{
  real alpha;                      // Effect of body wt in mediation model
  vector[P] beta;                  // Body weight coefs (group means)
  vector[P] theta;                 // Liver weight coefs (group means)
  vector[P] gamma;                 // Total effect on liver wt (group means)

  real<lower=0> sigma_beta;        // Variation between betas
  real<lower=0> sigma_theta;       // Variation between thetas
  real<lower=0> sigma_gamma;       // Variation between gammas

  vector<lower=0>[P] bw_var;       // Body weight within-group variation
  vector<lower=0>[P] lw_var;       // Liver weight within-group variation
  vector<lower=0>[P] lw_var_tot;   // Liver weight within-group variation

  real<lower=0> bw_var_sigma;      // Variation between bw_vars
  real<lower=0> lw_var_sigma;      // Variation between lw_vars
  real<lower=0> bw_var_tot_sigma;  // Variation between lw_var_tots
}

model{
  // priors
  sigma_beta ~ normal(0, 120);
  sigma_theta ~ normal(0, 10);
  sigma_gamma ~ normal(0, 120);

  alpha ~ normal(0, 10);
  beta ~ student_t(3, 0, sigma_beta);
  theta ~ student_t(3, 0, sigma_theta);
  gamma ~ student_t(3, 0, sigma_gamma);

  bw_var ~ normal(0, bw_var_sigma);
  lw_var ~ normal(0, lw_var_sigma);
  lw_var_tot ~ normal(0, bw_var_tot_sigma);

  bw_var_sigma ~ normal(0, 140);
  lw_var_sigma ~ normal(0, 10);
  bw_var_tot_sigma ~ normal(0, 140);

  // likelihood
  if(prior_only == 0){
    for(i in 1:N){
      bw[i] ~ normal(beta[grp[i]], bw_var[grp[i]]);
      lw[i] ~ normal(theta[grp[i]] + alpha*bw[i], lw_var[grp[i]]);
      
      // not necessary as can be calculated, but useful to check
      lw[i] ~ normal(gamma[grp[i]], lw_var_tot[grp[i]]);
    }
  }
}

generated quantities {
  //vector[N] log_lik;
  vector[N] bw_sim;   // Body weight
  vector[N] lw1_sim;  // Liver weight, adjusting for body weight
  vector[N] lw2_sim;  // Liver weight (no adjustment)

  // posterior predictive
  for (i in 1:N) {
    //log_lik[i] = normal_lpdf(Y[i] | beta[grp[i]], sigma);
    bw_sim[i] = normal_rng(beta[grp[i]], bw_var[grp[i]]);
    lw1_sim[i] = normal_rng(theta[grp[i]] + alpha*bw[i], lw_var[grp[i]]);
    lw2_sim[i] = normal_rng(gamma[grp[i]], lw_var_tot[grp[i]]);
  }
}
```
