## Supplementary material for "Determining organ weight toxicity with Bayesian causal models: Improving on the analysis of relative organ weights": Data and R code.: simulations.stan

```
data{
  int N;                    // Number of animals
}

generated quantities {
  //  real alpha;               
  // real<lower=0> bw_var;     
  // real<lower=0> lw_var;     
  vector[N] bw_sim;         // Body weight
  vector[N] lw_sim;         // Liver weight, adjusting for body weight

  real alpha = 0.0435;      // Effect of body wt in mediation model
  real bw_var = 16.2;       // Body weight within-group variation
  real lw_var = 0.66;       // Liver weight within-group variation
  
  // posterior predictive
  for (i in 1:N) { 
    bw_sim[i] = normal_rng(0, bw_var);
    lw_sim[i] = normal_rng(0 + alpha*bw_sim[i], lw_var);
  }
}
```
